# Spatial profiling identifies regionally distinct microenvironments and targetable immunosuppressive mechanisms in pediatric osteosarcoma pulmonary metastases

**DOI:** 10.1101/2025.01.22.631350

**Authors:** Jason Eigenbrood, Nathan Wong, Paul Mallory, Janice Pereira, Douglass W Morris, Jessica A Beck, James C Cronk, Carly M Sayers, Monica Mendez, Linus Kaiser, Julie Galindo, Jatinder Singh, Ashley Cardamone, Milind Pore, Michael Kelly, Amy K LeBlanc, Jennifer Cotter, Rosandra N Kaplan, Troy A McEachron

## Abstract

Osteosarcoma is the most common malignant bone tumor in young patients and remains a significant clinical challenge, particularly in the context of metastatic disease. Despite extensive documentation of genomic alterations in osteosarcoma, studies detailing the immunosuppressive mechanisms within the metastatic osteosarcoma microenvironment are lacking. Our objective was to characterize the spatial transcriptional landscape of metastatic osteosarcoma to reveal these immunosuppressive mechanisms and identify promising therapeutic targets. Here, we performed spatial transcriptional profiling on a cohort of osteosarcoma pulmonary metastases from pediatric patients. We reveal a conserved spatial gene expression pattern resembling a foreign body granuloma, characterized by peripheral inflammatory signaling, fibrocollagenous encapsulation, lymphocyte exclusion, and peritumoral macrophage accumulation. We also show that the intratumoral microenvironment of these lesions lack inflammatory signaling. Additionally, we identified CXCR4 as an actionable immunomodulatory target that bridges both the intratumoral and extratumoral microenvironments and highlights the spatial heterogeneity and complexity of this pathway. Collectively, this study reveals that metastatic osteosarcoma specimens are comprised of multiple regionally distinct immunosuppressive microenvironments.

## INTRODUCTION

Osteosarcoma, the predominant malignant bone tumor in pediatric, adolescent, and young adult patients, poses a significant clinical challenge as there has been little improvement in outcomes over the past four decades.^1–3^ The presence of metastasis is the most significant adverse prognostic factor.^1,2^ Those patients diagnosed with metastatic osteosarcoma face notably inferior outcomes, with a median overall survival of approximately 20 months and 5-year survival of < 30%, in stark contrast to individuals with localized disease who have a 70% 5-year survival.^1,4^ The current standard of care involves surgical intervention and a high-dose multiagent chemotherapy regimen and is associated with significant surgical morbidity and systemic toxicities.^5^ Considering the discouraging outcomes for patients with metastatic osteosarcoma, this disease remains a pressing unmet medical need.^6^

While the genomic alterations driving this disease have been extensively documented, there remains a scarcity of studies investigating the spatial molecular and cellular composition of the osteosarcoma microenvironments and the immunosuppressive mechanisms operating within them. In the context of osteosarcoma pulmonary metastases, in which malignant bone is growing in a soft, well-oxygenated ectopic site, the microenvironmental features necessary to support metastatic malignant bone growth are largely unknown. Furthermore, the unique aspects of these cellular communities within the metastatic microenvironment have not been completely elucidated and therapeutic approaches targeting the tumor and microenvironmental crosstalk have yet to be realized for metastatic osteosarcoma.

Recent single-cell RNA sequencing studies have offered valuable insights into the cellular composition of osteosarcomas, primarily focusing on primary disease.^7–9^ However, whether and where these specific cell subsets reside within pulmonary metastases and how their spatial distribution contributes to the overall structure and function of the metastatic microenvironment cannot be assessed by single cell techniques. To understand the transcriptional architecture of metastatic osteosarcoma, identify regional gene expression programs, and gain insight into the immunosuppressive mechanisms associated with metastatic disease, we performed spatial transcriptional profiling on nine osteosarcoma pulmonary metastases derived from seven patients. Our data reveal that the spatial gene expression landscape of pulmonary metastases of osteosarcoma is remarkably conserved across specimens. We also show that the intratumoral microenvironment is hallmarked by limited pro-inflammatory chemokine/cytokine signaling and abundant TGFβ signaling. Lastly, by exploring spatially distinct drug-gene interactions, we identified CXCR4 as a clinically relevant therapeutic target for metastatic osteosarcoma.

## MATERIALS AND METHODS

### Specimen selection and quality assessment

Archived formalin-fixed paraffin-embedded (FFPE) specimens from pediatric osteosarcoma patients with clinically confirmed metastatic disease were obtained from the Department of Pathology at Children’s Hospital Los Angeles under IRB protocol CCI12-00224. Serial 5μm-thick sections were cut from each block. A single slide was stained using hematoxylin and eosin and subsequently used to evaluate the presence of both tumor and surrounding non-malignant tissue as well as specimen integrity of each slide. Specimens with substantial necrosis, hemorrhage, tissue tears, and/or folds were not used for Visium profiling. The Visium CytAssist Spatial Gene Expression assay relies on hybridization of RNA-binding probes. To ensure the tissues of interest were capable of hybridizing RNA-binding probes we used the RNAscope 2.5 Assay (ACD Bio) using probes against GAPDH (ACD Bio, #310321-C2) according to the manufacturers protocol. Specimens demonstrating a lack of RNAscope signal were considered unsuitable for Visium profiling and excluded. In total, 9 specimens surpassed our quality control criteria and were used for spatial profiling.

### Survival analysis

Kaplan-Meier survival curves were generated using data from the “RNAseq_Landscape_Manuscript” project in the online clinomics database (cliniomics.ccr.cancer.gov) hosted by the Oncogenomics Section of the Genetics Branch of the National Cancer Institute. Significance threshold was determined using p-value minimization.

### Spatial transcriptional profiling of FFPE specimen using the Visium CytAssist platform

Tissues were sectioned onto positively charged glass slides and destained and decrosslinked according to the 10X Genomics demonstrated protocol. Probe hybridization and all subsequent steps were performed as per 10x Genomics user guide for Visium CytAssist Spatial Gene Expression. In short, probe hybridization with human whole transcriptome probe set v2 was performed overnight on the prepared tissue sections with all samples receiving the same hybridization time. Post hybridization washes were followed by probe ligation, and ligated probes were released and transferred onto Visium spatially barcoded slides using the CytAssist platform, targeting regions of interest within 11x11mm capture areas. Subsequently, probe extension was carried out and the ligation product was eluted, and a pre-amplification step performed. A subset of the pre-amplification product was used for library construction after quantification by qPCR. Quality and concentration of the resulting sequencing libraries were checked using BioAnalyzer 2100 instrument (Agilent Technologies). The Visium libraries were sequenced on an Illumina NovaSeq 6000 sequencer with the following run parameters: 28bp (Read1), 10bp (Index1), 10bp (Index2), 50bp (Read2). Multiple sequencing runs were performed to target a depth of 50,000 reads per tissue-covered spatial barcode, with samples multiplexed across sequencing runs when possible. Illumina base calling used RTA 3.4.4. 10X Genomics Visium sequencing data demultiplexing was performed using SpaceRanger v2.0.0 (Bcl2fastq 2.20.0), and alignment was performed using SpaceRanger v2.0.0 (STAR2.7.2a). Sequenced reads were aligned to the 10X Genomics provided human reference sequence (refdata-gex-GRCh38-2020-A) and probe set (Visium Human Transcriptome Probe Set v2.0).

### Processing and cleaning the Visium dataset

All computational tools used in this study are published packages that are publicly available. All analyses were performed using R (v4.3.0) unless otherwise specified. SpotClean was used to correct for spot swapping.^10^ A maximum of 30 iterations and a candidate radius of 20 was used to correct for spot swapping. The decontaminated object from SpotClean was then imported into Seurat (v4.2 – v5.1) for further downstream processing. The Seurat object was filtered to exclude spots with extremely low numbers of counts and gene as well as those with extremely high number of counts and gene, based on histograms displaying number of counts per spot and number of expressed genes per spot. Each specimen was normalized using the “SCTransform” function in Seurat. Marker genes were identified for each cluster using the “FindAllMarkers” function with a log fold change threshold of 0.5.

### Generation of the intratumoral osteosarcoma gene signature

To resolve the heterogeneity of the intratumoral microenvironment and infer the relative cellular proportions within each spot without a robust metastatic osteosarcoma single cell RNA sequencing reference, we applied STDeconvolve to our dataset.^11^ Other than increasing the maximum number of potential cell types, or “K,” from 9 to 20, the default parameters were used for this analysis. Up to 18 purported deconvolved cell types (referred to as “topics”), were identified per sample. For each potential topic, the top 100 elevated gene markers were used to generate correlation scores between all topics within and across samples. Correlation analysis was performed on the STDeconvolve outputs to identify the most highly ranked genes that map to the intratumoral microenvironment in at least 6 of the 9 specimens. To this list, we added a manually curated list of marker genes that recurrently mapped to intratumoral clusters and excluded genes expressed in tissue resident fibroblast subsets using the Human Lung Cell Atlas as a reference.^12^ This resulted in a 23 gene Osteosarcoma Signature (**Supplemental Table 1**).

### Visualization of spatial transcriptomics data

Uniform Manifold Approximation and Projection (UMAP) plots and spatial cluster maps of the cluster assignments were made using the Seurat functions “DimPlot” and “SpatialDimPlot”, respectively. UMAP plots and spatial maps showing genes and/or gene signatures were made using the Seurat functions “FeaturePlot” and “SpatialFeaturePlot” respectively. To improve the visualization of the data, the “image.alpha” was set to 0, the “min.cutoff” was set to “q10”, and the spot size was adjusted for each specimen. Clustered dot plots were generated using scCustomize (https://samuel-marsh.github.io/scCustomize/). The “corrplot” was used to generate hierarchically clustered correlation matrices. Heatmaps were generated using Morpheus (https://software.broadinstitute.org/morpheus).

### Gene set enrichment analysis

The differentially enriched gene sets in each cluster were determined using Escape.^13^ Enrichment was performed using the “enrichIt” function using the Hallmarks, Reactome, and Biological Processes gene set libraries from MSigDB. We utilized the “UCell” method with a “maxRank” set to 2000. The “getSignificance” function was used to determine the significantly enriched gene sets in each cluster by ANOVA.

### Scoring metagene signatures

The UCell package was used to assign cell type specific metagene signatures based on single cell RNA sequencing reference datasets.^14^ Gene signatures from the Human Lung Cell Atlas and the Human PBMC reference datasets were obtained from the Azimuth database.^12,15^ The metagene signature scores were added to the Seurat object metadata using the “AddModuleScore_UCell” function. Cell type specific gene signatures are listed in **Supplemental Table 1**.

### Inferred cytokine activity

Cytokine signaling activity predictions were performed using the web-based instance of CytoSig.^16^ First, the average gene expression for each transcriptional cluster was calculated for individual Seurat objects with the function “AverageExpression”. This output was uploaded to https://cytosig.ccr.cancer.gov/profiler/task_upload/ to predict the cytokine signaling activity in each cluster. The computed cytokine activity scores were then added to the metadata of each Seurat object. The “MapCytosig” function was used to spatially map the cytokine activity scores, thereby revealing the inferred signaling activity of a given cytokine within each transcriptional cluster.

### Ligand-receptor interactions

The CellChat package was used to elucidate spatially constrained ligand-receptor interactions.^17^ Default parameters were used to create The CellChat object was created using the the following exceptions: spot.size = 55; contact.range = 100, interaction.range = 250, and scale.distance = 0.01.

### Generation of the integrated spatial dataset

Harmony was used to integrate the individual Seurat objects using the following values: “theta=2, res=0.3, dims=15”.^18^ Marker genes from each integrated cluster were uploaded to the EnrichR website (https://maayanlab.cloud/Enrichr/).^19–21^ The “BioPlanet 2019”, “KEGG 2021”, “Reactome 2022”, and “WikiPathways 2023” libraries were used to identify differentially enriched pathways and processes in each integrated cluster.

### Drug-gene interaction analysis

Spatial drug-gene interactions were determined using the “IDG Drug Targets 2022’ library in EnrichR with the marker genes from each Integrated Cluster serving as input. The enrichment score results of the drug-gene interaction are presented as the combined score. Histograms were generated using Prism 10 (GraphPad).

### Imaging mass cytometry (IMC)

Metal-labeled antibodies previously validated for immunohistochemistry and IMC were purchased from Standard BioTools (**Supplemental Table 2**). Hematoxylin and eosin stained FFPE sections were used to identify regions of interest (ROIs) for laser ablation on the Hyperion Imaging System (Standard BioTools). FFPE sections were deparaffinized, rehydrated through graded ethanols, and subjected to heat-mediated antigen retrieval in Tris-EDTA buffer (pH 9) at 95 °C for 30 minutes. The tissue sections were blocked with 3%BSA in PBS, incubated with the metal-labeled antibody cocktail overnight at 4^0^C, then counterstained with Cell-ID DNA intercalator for 30 minutes. Slides were washed in PBS, dipped in deionized water to remove the salt traces, and dried using pressurized air. Prior to the sample acquisition, the Hyperion Imaging System was autotuned using 3-element tuning slide as described in the Hyperion user guide. Each ablated ROI measured 1.5 mm^2^. The raw IMC images were denoised using DIMR and Deep-SNiF from the IMC-Denoise toolset to reduce image artifacts.^22^ The denoised images were then preprocessed using the IMC Segmentation Pipeline (https://github.com/BodenmillerGroup/ImcSegmentationPipeline) created by the Bodenmiller Laboratory. A pretrained IMC nuclei detection convolutional neural network from was used to generate cell segmentation masks (Visiopharm). Single-cell mean marker intensities and spatial features were extracted from the denoised images using the segmentations masks and were subsequently used for downstream analysis and visualization following the Bodenmiller pipeline.^23^

### Cell lines

Patient derived osteosarcoma cell lines were provided by Dr. Alejandro Sweet-Cordero.^24^ All cell lines were confirmed free of mycoplasma contamination. Early passage stocks of each line were used to collect conditioned media. Conditioned media was collected as follows: 1x10^6^ cells were plated in in DMEM (Invitrogen) + 10% Bovine Growth Serum (Cytiva). After 24-hours of growth, the cells were washed and fresh growth media was added for 24-hours, at which point the conditioned media was collected, cleared of debris by centrifugation, and stored at -30C until use. The conditioned media was harvested from three independent biological replicates.

### Cytokine profiling arrays

Cytokine arrays were performed using the Human XL Cytokine Array kit (R&D Systems #ARY022B) according to the manufacturer’s instructions. 200μg of human metastatic osteosarcoma protein lysate was used per array. For each cell line, equal volumes of conditioned media from each biological replicate was pooled and then added to the array. Quick Spots software (Ideal Eyes) was used to analyze the data. Each array was background subtracted and normalized to the average of the positive control spots.

### Immunohistochemistry

Immunohistochemistry was performed as previously described.^25^ Briefly, FFPE sections were deparaffinized, rehydrated, and subjected to heat mediated antigen retrieval in citrate buffer. Specimens were then blocked with horse serum incubated with a rabbit monoclonal antibody against SMAD4 (Cell Signaling Technologies #85061) or a rabbit monoclonal antibody against p65 (Cell Signaling Technologies #8242) overnight at 4C. Color development was achieved by incubating the slides with a peroxidase conjugated anti-rabbit secondary antibody (Vector Laboratories, #MP-7401) followed by 3, 3’-diaminobenzidine (Vector Laboratories, SK-4105). Images were captured on an AxioScan 7 slide scanner with a 20X objective (Zeiss).

### Trichrome staining

Trichrome staining was performed a commercial trichrome stain kit (Abcam, #ab150686) according to the manufacturers recommended protocol.

### Data availability

Deposition of the expression data in the Gene Expression Omnibus (GEO) is currently in progress. Once completed, the expression data will be made publicly available, and the manuscript will be updated to include the associated accession number.

## RESULTS

### Resolving the spatial transcriptome of individual osteosarcoma pulmonary metastases

To elucidate regional gene expression profiles and transcriptional programs in metastatic osteosarcoma specimens, we performed spatial transcriptional profiling using the Visium CytAssist platform. In total, nine pathologically confirmed FFPE osteosarcoma pulmonary metastases specimens obtained from seven individual pediatric patients were assayed. Of these, seven specimens were obtained from the lung parenchyma while two specimens were metastatic hilar lymph nodes. Specimens were selected on the basis of containing sufficient adjacent non-malignant tissue to comprehensively investigate the tumor-non-tumor interface. Each specimen was reviewed and annotated by a pathologist with significant experience in sarcoma histopathology (JB) to identify the malignant regions. Based on this pathological evaluation, each annotated specimen was divided into intratumoral and extratumoral regions (**Figure 1A**). With little exception, unsupervised Leiden clustering of the gene expression data partitioned the spots from each specimen into subclusters localized within the tumor core (intratumoral) and outside of the tumor core (extratumoral) as demonstrated by both spatial maps and UMAP plots (Figure 1B-C). Despite the independent analysis and clustering of each specimen, the existence of a well-defined circumscribing peritumoral microenvironment was a consistent finding across the specimens, with the exception of fibroblastic osteosarcoma specimen OS-7 (**Figure 1C**). The *in situ* distribution and localization of these different transcriptional clusters indicates that metastatic osteosarcoma specimens are well structured tissues with spatially and transcriptionally distinct microenvironments.

**Figure 1:**
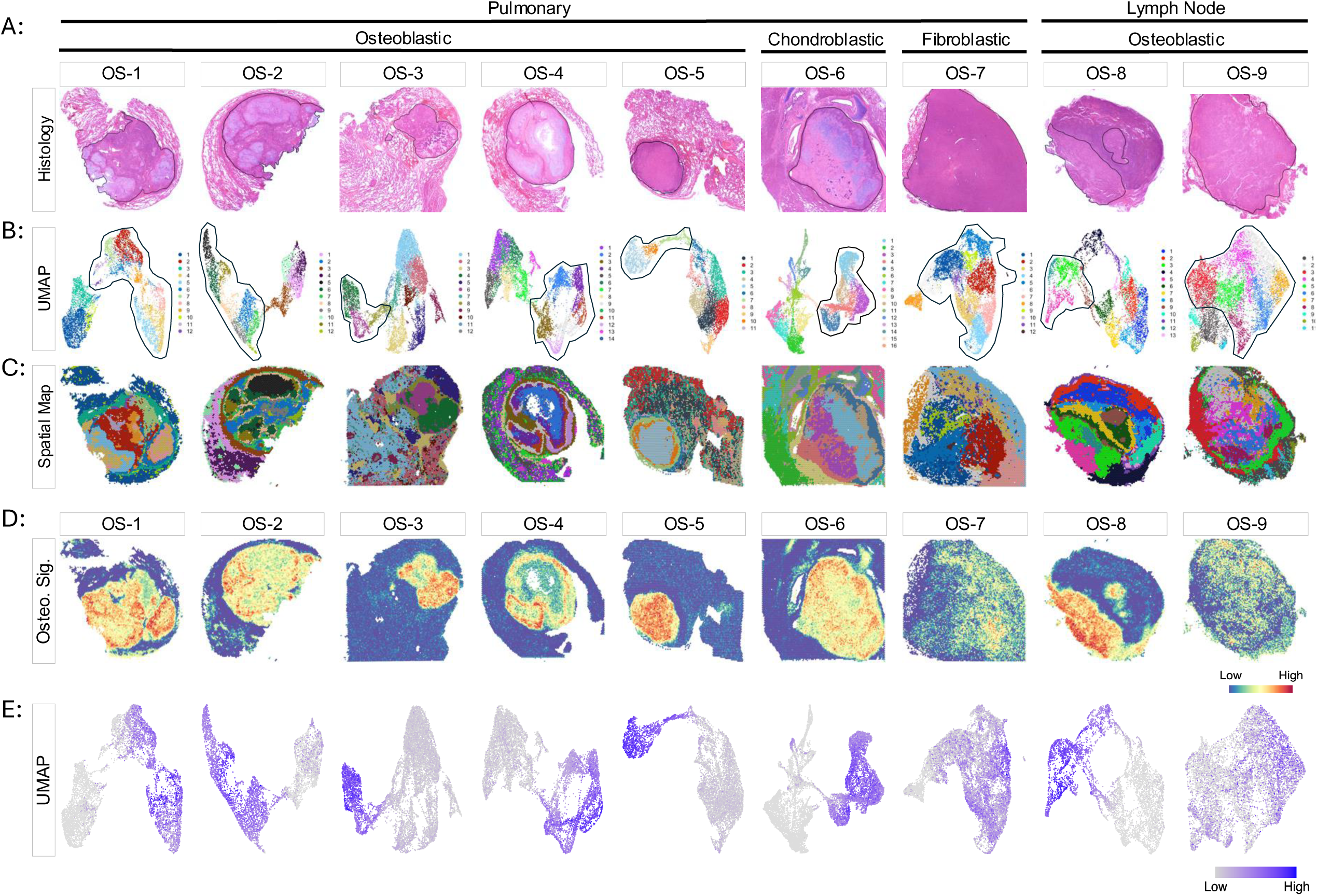
The transcriptional architecture of metastatic osteosarcoma. (A) Annotated hematoxylin and eosin-stained histological images. Malignant regions of each tissue are outlined in black based on expert pathological assessment. Disease site and histology are indicated above. (B) UMAP plots of the transcriptionally distinct clusters identified in each specimen. Separate color palettes were used for each specimen to denote the independent clustering analysis of each specimen. The clusters corresponding to the malignant regions of each tissue are outlined in black. (C) Spatial cluster maps of each specimen showing the *in-situ* distribution of transcriptional clusters displayed on the above UMAP plots projected on to the histological images. (D) Spatial mapping and (E) UMAP plots of the Osteosarcoma Signature (Osteo. Sig.), highlighting the intratumoral osteosarcoma microenvironment in each specimen.

### Defining the transcriptional signature of the intratumoral metastatic osteosarcoma microenvironment

Osteosarcomas are histologically heterogeneous tumors with malignant osteoblastic, chondroblastic, and fibroblastic features of varying abundance frequently observed within a single specimen.^26^ The 55µm spot size of Visium assay can capture gene expression profiles from multiple cell types in any given spot. Additionally, osteosarcoma cells express genes that overlap with non-malignant mesenchymal lineage cells and myeloid cell subsets. For these reasons, we were unable to confidently infer individual cellular identities within the tumor core. As an alternative, we created a generalized gene expression signature that delineates the intratumoral metastatic osteosarcoma microenvironment from the adjacent non-malignant regions of the tissue (**Supplemental Table 1**). Spatial mapping of our “Osteosarcoma Signature” mirrored the expert pathological annotations of the H&E images (**Figure 1A, D-E**). In specimens OS-7 and OS-9, this gene signature helped to discern regions of high and low tumor content that were histologically annotated as tumor (**Figure 1A, D-E**).

### Spatial distribution and abundance of myeloid lineage cells

Immunohistochemical analysis and bulk gene expression profiling has previously demonstrated that T-cells are excluded from infiltrating osteosarcoma pulmonary metastases, whereas myeloid cells are a prominent cellular component of the intratumoral metastatic osteosarcoma microenvironment.^25,27^ Myeloid cells demonstrate microenvironment-dependent functional plasticity, capable of both hindering or enhancing tumor progression, therefore gene signatures used to broadly identify myeloid subsets cannot be assumed to imply function.^28–31^ Analysis of individual genes associated with macrophage function uncovered robust spatial gene expression patterns across all compartments and specimens. In each specimen, a subset of spatial clusters segregated independently from the other extratumoral and intratumoral spatial clusters, indicating a markedly different transcriptional profile (**Supplemental Figure 1**). Indeed, within each of these distinct transcriptionally active clusters, a multitude of phenotypic and functional macrophage genes were expressed, including *C1QA, CCL18, CD68, MARCO, MRC1* (encoding CD206)*, MSR1,* and *TREM1* (**Supplemental Figure 1**). The genes in these corresponding clusters have been associated with the suppression of adaptive immune responses.^32–35^ The M2 macrophage markers *CSF1R, PTGES2,* and *FOLR2* were primarily expressed within the tumor core whereas *MARCO, C1Q,* and *S100A9* were expressed in the extratumoral regions of the tissues (**Supplemental Figure 1**). Together, these data suggest that phenotypically and functionally distinct immunosuppressive myeloid subsets occupy specific intratumoral and extratumoral microenvironments.^35,36^

### Evidence of a peritumoral foreign body-like response

Given the localized enrichment of numerous immune cell signatures and the assorted repertoire of both phenotypic and functional macrophage-associated genes within the peritumoral microenvironment, we decided to investigate this region in greater depth. Upon careful histopathological examination of the tissues, we identified peritumoral multinucleated giant cells resemblant of Langhans giant cells (**Figure 2A**).^37^ The appearance of these cells is consistent with a foreign body response, an ordered series of events that aims to isolate biological and/or synthetic material that the immune system has not been able to clear.^38,39^ The final stage of the foreign body response is the encapsulation of the material in a fibrocollagenous matrix. Trichrome staining of the specimens revealed a pronounced deposition of collagen circumscribing the tumor core in both the lung and lymph node metastases (**Figure 2B-C**). Spatially resolved gene set enrichment analysis using the MSigDB gene ontology dataset revealed that the Defense Response signature, a transcriptional surrogate for the response to a foreign body or injury, was enriched in the extratumoral microenvironment surrounding the tumor core (**Figure 2D**). Spearman correlation analysis was performed to investigate the relationships between the Defense Response signature and the various inferred cell type-specific signatures. This analysis showed significant inverse relationships between the Osteosarcoma and Defense Response signatures (**Figure 2D-E**).

**Figure 2:**
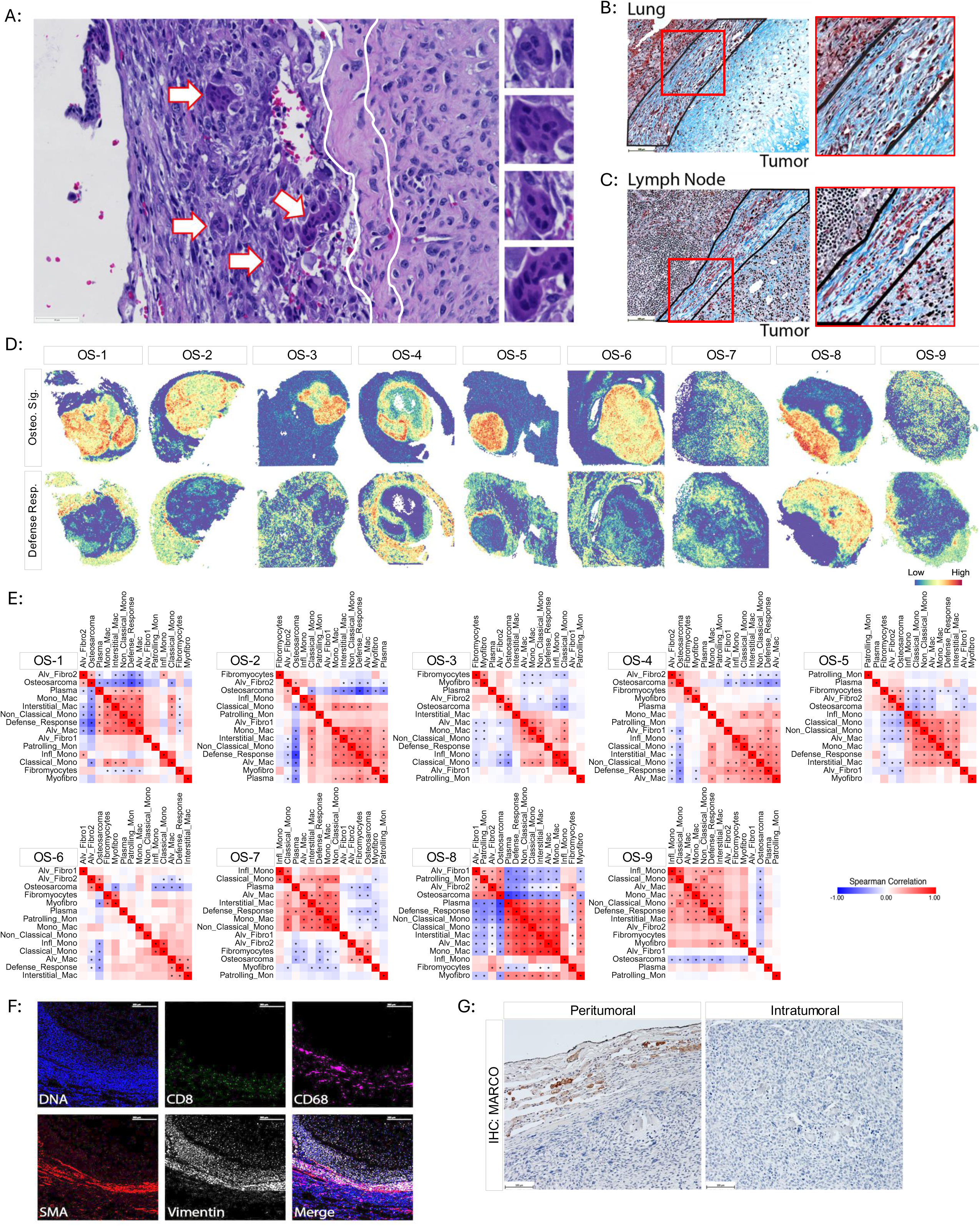
Evidence a foreign body response and fibrocollagenous encapsulation in pulmonary metastatic lesions. (A) Representative hematoxylin and eosin-stained metastatic osteosarcoma specimen highlighting the appearance of peritumoral multinucleated giant cells resemblant of Langhans giant cells (arrows) adjacent to the tumor core/intratumoral microenvironment. Right: Enlarged images of multinucleated cells identified in the image to the left. White line outlines the region of circumscribing collagen deposition separating the peritumoral environment from the tumor core/intratumoral microenvironment. (B, C) Representative trichrome staining showing circumscribing deposition of collagen (outlined in black) separating the tumor core/intratumoral microenvironment from the non-malignant (B) lung stroma and (C) lymph node tissue. Right: Enlarged images of the insets. (D) Spatial mapping of the Defense Response (Defense Resp.) gene signature. Spatial plots highlighting Osteosarcoma Signature (Osteo. Sig.) are shown for reference. (E). Spearman correlation matrices for each specimen comparing the Osteosarcoma Signature, Defense Response signature, and signatures of different monocyte, macrophage, and fibroblast subsets. Asterisks indicate a significant correlation with p ≤ 0.05. (F) Imaging mass cytometry of the tumor interface. Individual channels were pseudocolored for visualization. Scale bar = 300µm. (G) Representative IHC staining of MARCO. Both images taken from different regions of the same specimen. Scale bar = 100µm.

We then confirmed these findings at the protein level using IMC. Antibodies against CD8 and CD68 were used to detect cytotoxic T-cells and macrophages, respectively. SMA is expressed by activated fibroblasts and myofibroblasts in various pathological contexts, including foreign body reactions.^40^ Vimentin is highly expressed in osteosarcoma cells and is also expressed in immune cells, including myeloid cells, and non-malignant mesenchymal cells, and thus served as a positive staining control.^41–43^ We observed that CD68^+^ macrophages, SMA^+^ fibroblasts, and CD8^+^ T-cells all localized to the peritumoral region in a concentric fashion (**Figure 2F**). These results mirror those obtained by spatial transcriptional profiling (**Figure 2D**). Together, these data suggest that the presence of metastatic disease in both the lung parenchyma and hilar lymph nodes provokes a foreign body-like response.

### MARCO expressing macrophages colocalize with the peritumoral foreign body-like response

Next, we sought to identify candidate cell subsets associated with the peritumoral foreign body-like response. In each specimen, the Alveolar Macrophage, Interstitial Macrophage, and Monocyte-derived Macrophage gene signatures positively correlated and colocalized with the Defense Response signature (**Figure 2E**). To further confirm these findings, we performed immunohistochemistry to detect MARCO, a scavenger receptor expressed at the mRNA level in both alveolar and interstitial macrophages.^44^ MARCO^+^ macrophages were observed in the peritumoral microenvironment and absent in the intratumoral microenvironment (**Figure 2G**). Both recruited, and tissue-resident macrophages have been shown to orchestrate pulmonary fibrosis and foreign body responses.^45–47^ Our data associates MARCO^+^ macrophages with the peritumoral foreign body-like response in metastatic osteosarcoma and may represent an important therapeutic target.

### Regionally restricted chemokine, cytokine, and growth factor activity

Next, we examined cellular communication networks to infer signaling activity within and between different regions in each specimen. The CytoSig package was used to quantify chemokine and cytokine activity within individual spatial clusters. Spearman correlation analysis of the CytoSig scores followed by unsupervised hierarchical clustering consistently partitioned each specimen into two independent groups with the intratumoral spatial clusters comprising one group and the extratumoral spatial clusters comprising the other (**Figure 3A**). Both IFNγ and TNFα are important inflammatory mediators in the tumor microenvironment. Spatial mapping of IFNγ and TNFα activity in individual specimens showed clear regional distinction with signaling activity exclusively confined to the extratumoral microenvironment (**Figure 3B**). Differential enrichment analysis of the aggregate CytoSig activity scores across all specimens revealed that inflammatory chemokine and cytokine activity preferentially localized to the extratumoral regions of the tissue whereas growth factor activity localized to the intratumoral microenvironment (**Figure 3C**).

**Figure 3:**
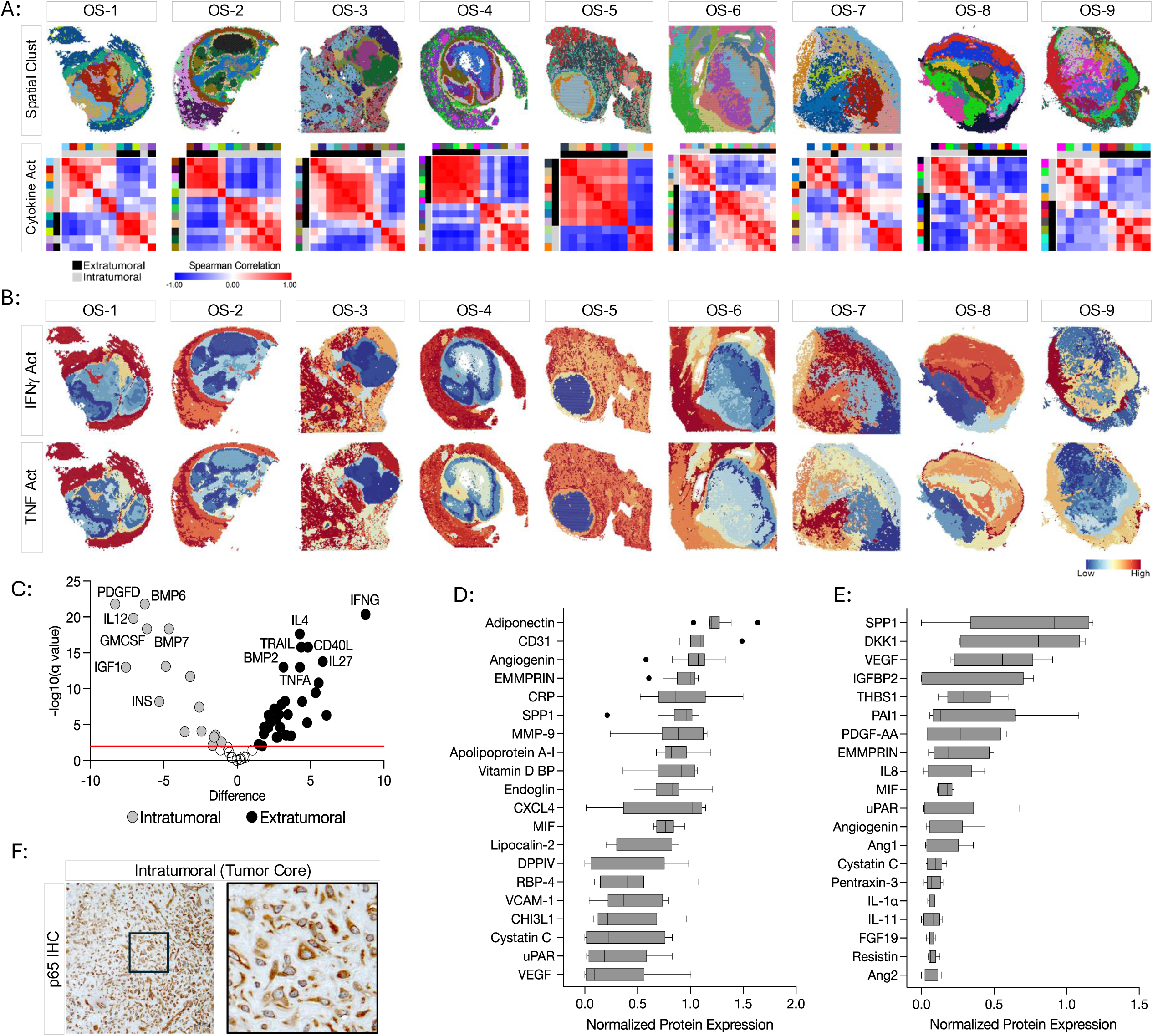
Differential cytokine signaling activity between the intratumoral and the extratumoral microenvironments. (A) Spatial cluster maps (top row) are shown for reference. Spearman correlation matrix of CytoSig Scores in each spatially defined cluster (bottom row). The top color bar above each heatmap corresponds to the spatial clusters displayed on the above spatial cluster maps. Separate color palettes denote the independent clustering analysis of each specimen. Spatial clusters were localized to the extratumoral (black) or intratumoral (gray) microenvironments based on the pathological annotations and the Osteosarcoma Signature and depicted in the second color bar. (B) Spatial plots illustrating the extratumoral localization of IFNγ and TNFα activity inferred by CytoSig. (C) Volcano plot of the aggregated differentially enriched CytoSig Scores between the extratumoral (black) and intratumoral (gray) spatial clusters. Horizontal red line indicates significance threshold (q = 0.001). (D-E) Top 20 proteins identified from cytokine antibody array profiling in (D) tissue homogenates from an independent cohort of frozen metastatic osteosarcoma specimens, and (E) conditioned media from patient derived cell lines. (F) Representative IHC staining of p65/NFκB. Scale bar = 50µm. Right: Enlarged image of the inset.

### The intratumoral microenvironment of metastatic osteosarcoma is an inflammatory desert

Our data shows that the intratumoral microenvironment lacks transcriptional programs indicative of inflammatory cytokine activity with IFNγ and TNFα signaling localized to the extratumoral regions (**Figure 3A-C**). To validate these findings at the protein level, homogenized tissue lysates from an independent cohort of human osteosarcoma pulmonary metastases were profiled using highly multiplexed antibody arrays to detect 105 individual chemokines/cytokines. Pathology reports confirmed that each specimen was of high tumor cellularity and viability. Conditioned media from patient derived cell lines was also profiled to assess the tumor cell intrinsic chemokine/cytokine profiles and to help delineate chemokines/cytokines that may be derived from tumor cells versus those from secreted by other cells present within the intratumoral microenvironment of the patient specimens. In both the tissue lysates and conditioned media, most of the inflammatory chemokines and cytokines assayed were below the detection threshold, with little or no protein expression (**Figure 3D-E, Supplemental Figure 2**). SPP1, VEGFA, EMMPRIN, MIF, uPAR, Angiogenin, and Cystatin C were among the most highly expressed analytes in both patient specimens and in cell line conditioned media, demonstrating that the osteosarcoma cells themselves are a cellular source of these molecules (**Figure 3D-E**). Despite its computationally inferred signaling activity within the intratumoral microenvironment, IL12 p70 protein was not detectable by cytokine array (**Figure 3C-E, Supplemental Figure 2**).

NFkB signaling, assessed by nuclear translocation of transcription factors p50, p52, p65, and/or RelB, is active in response to various stimuli including numerous pro-inflammatory chemokines and cytokines.^48^ As a surrogate marker for inflammatory signaling, we performed immunohistochemistry using an anti-p65 antibody. While cytoplasmic immunoreactivity was abundant, we did not observe nuclear p65 within the intratumoral microenvironment (**Figure 3F**). We interpreted these data to indicate a lack of robust intratumoral inflammatory signaling, supporting the minimal levels of computationally inferred cytokine activity within the intratumoral microenvironment. These data support bulk gene expression profiling data from an independent cohort of specimens showing that, in comparison to primary tumors, metastatic osteosarcoma specimens express a limited number of genes encoding inflammatory mediators (**Supplemental Figure 3**). Together, these data demonstrate minimal transcriptional and proteomic inflammatory chemokine/cytokine expression in the intratumoral microenvironment.

### Inflammatory signaling is enriched in the peritumoral microenvironment

Interactome analysis was performed to identify regionally constrained inflammatory ligand-receptor interactions. Specifically, we investigated the ligand-receptor utilization of TNFα, CXCL-, CCL-, and IL-pathways within and between the spatial clusters. Repeatedly, the spatial clusters corresponding to the intratumoral microenvironment were among the least communicative, whereas the peritumoral clusters were highly communicative (**Figure 4**). These data demonstrate that inflammatory chemokine and cytokine signaling activity in metastatic osteosarcoma is highly compartmentalized and regionally distinct and that the peritumoral microenvironment is a hub of inflammatory signaling.

**Figure 4:**
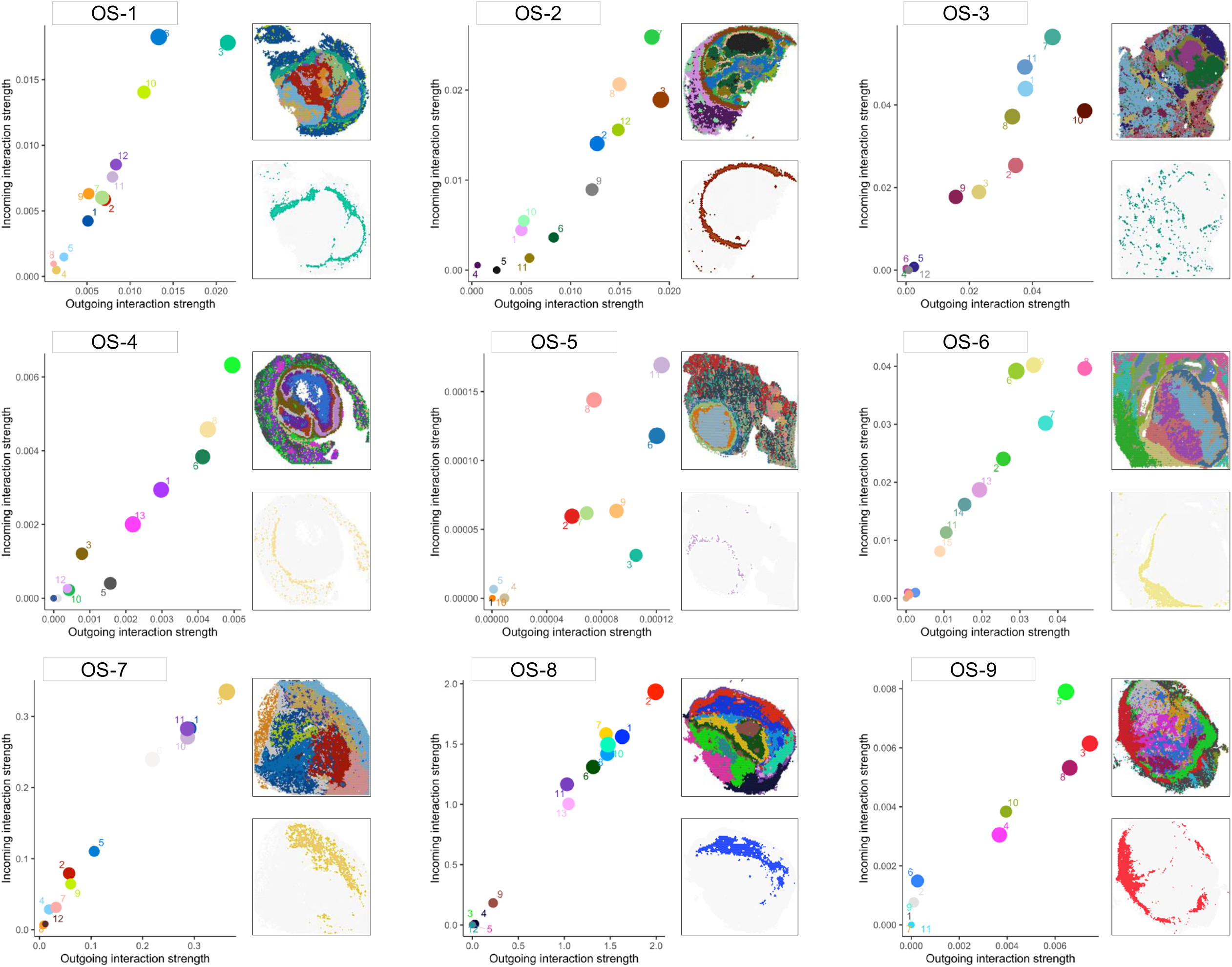
Identification of the most communicative spatial clusters based on ligand-receptor analysis. Scatter plots quantifying the combined outgoing (x-axis) versus incoming (y-axis) ligand-receptor interaction strength of “TNFA”, “CXCL-“, “CCL-“, and “IL-“ pathways. Dots represent individual spatial clusters and are colored according to spatial plots shown in the insets. Dots in the top right quadrant indicate the most active/dominant communicating regions in the tissue. Spatial maps displaying highly communicative clusters are shown to the right of each graph to visualize the cluster location within the tissue.

### Widespread and heterogeneous signaling activity by TGFβ family members

In addition to its immunoregulatory and profibrotic functions, TGFβ signaling is necessary for both physiological as well as pathological bone growth.^49^ We examined the regional distribution of TGFβ signaling activity and extended this inquiry to include multiple members of the TGFβ family (**Figure 5A**). Analysis of individual specimens showed heterogeneous and widespread TGFβ activity throughout all regions of the tissues with BMP6 and BMP7 activity primarily restricted to the intratumoral microenvironment (**Figure 5B-C**). Ligand-receptor analysis confirmed the presence of a complex and heterogeneous TGFβ interactome in each specimen (**Figure 5D**). Canonical TGFβ signaling results in nuclear translocation of SMAD4 (**Figure 5A**). Nuclear SMAD4 immunoreactivity was detected in the intratumoral microenvironment across all specimens (**Figure 5E**), confirming activation of the canonical SMAD4-dependent TGFβ signaling pathway within the intratumoral microenvironment.

**Figure 5:**
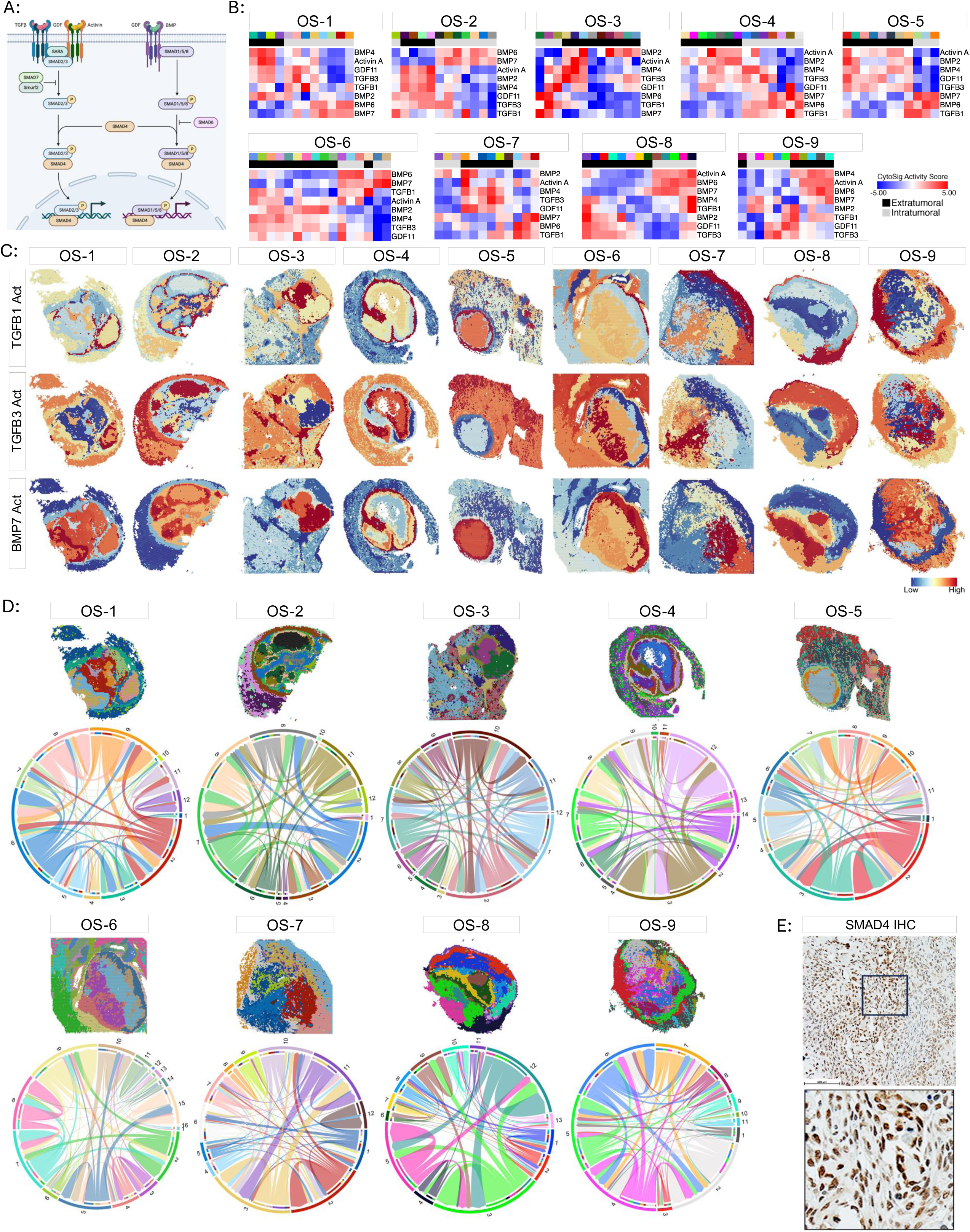
Widespread and heterogeneous signaling of TGFβ family members. (A) Illustration of canonical TGFβ family member signaling. Illustration made using Biorender. (B) Heatmaps of Activin A, BMP2, BMP4, BMP6, BMP7, GDF11, TGFβ1, and TGFβ3 CytoSig Scores. The color bar above each heatmap corresponds to the spatial clusters displayed on the associated spatial cluster maps. Separate color palettes denote the independent clustering analysis of each specimen. Spatial clusters were localized to the extratumoral (black) or intratumoral (gray) microenvironments based on the pathological annotations and the Osteosarcoma Signature and depicted in the second color bar. (C) Spatial plots illustrating the localization of TGFβ1 (top row), TGFβ3 (middle row), and BMP7 (bottom row) activity inferred by CytoSig. (D) Spatially resolved TGFβ-family ligand-receptor interactome within and between spatially defined clusters. Spatial cluster maps are shown for reference. Interactome chord diagram is colored according to the spatial clusters displayed on the above spatial cluster maps. (E) Representative IHC staining of SMAD4 in the intratumoral microenvironment. Scale bar = 100µm. Below: Enlarged image of the inset.

### Data integration identifies a conserved transcriptional architecture across specimens

Independent analysis of each specimen revealed a highly similar spatial distribution of transcriptional programs across the cohort (**Figures 1-5, Supplemental Figures 1, 3**). Next, we integrated our dataset to identify shared region-specific transcriptional programs and biological processes. Unsupervised hierarchical clustering resulted in 10 integrated clusters, termed “IC’s”, each localizing to similar regions across the different specimens (**Figure 6A-C**). The marker genes from each integrated cluster were used to identify conserved region-specific biological pathways (**Figure 6C**). Consistent with our results obtained by analyzing each specimen independently, pathways associated with inflammation and immune activity localized to the extratumoral integrated clusters (**Figure 6C**). In contrast, processes associated with cell cycle and ossification were enriched in the intratumoral IC’s (**Figure 6C**). These findings demonstrate that region-specific gene expression profiles and biological processes are conserved among the metastatic osteosarcoma specimens in this study.

**Figure 6:**
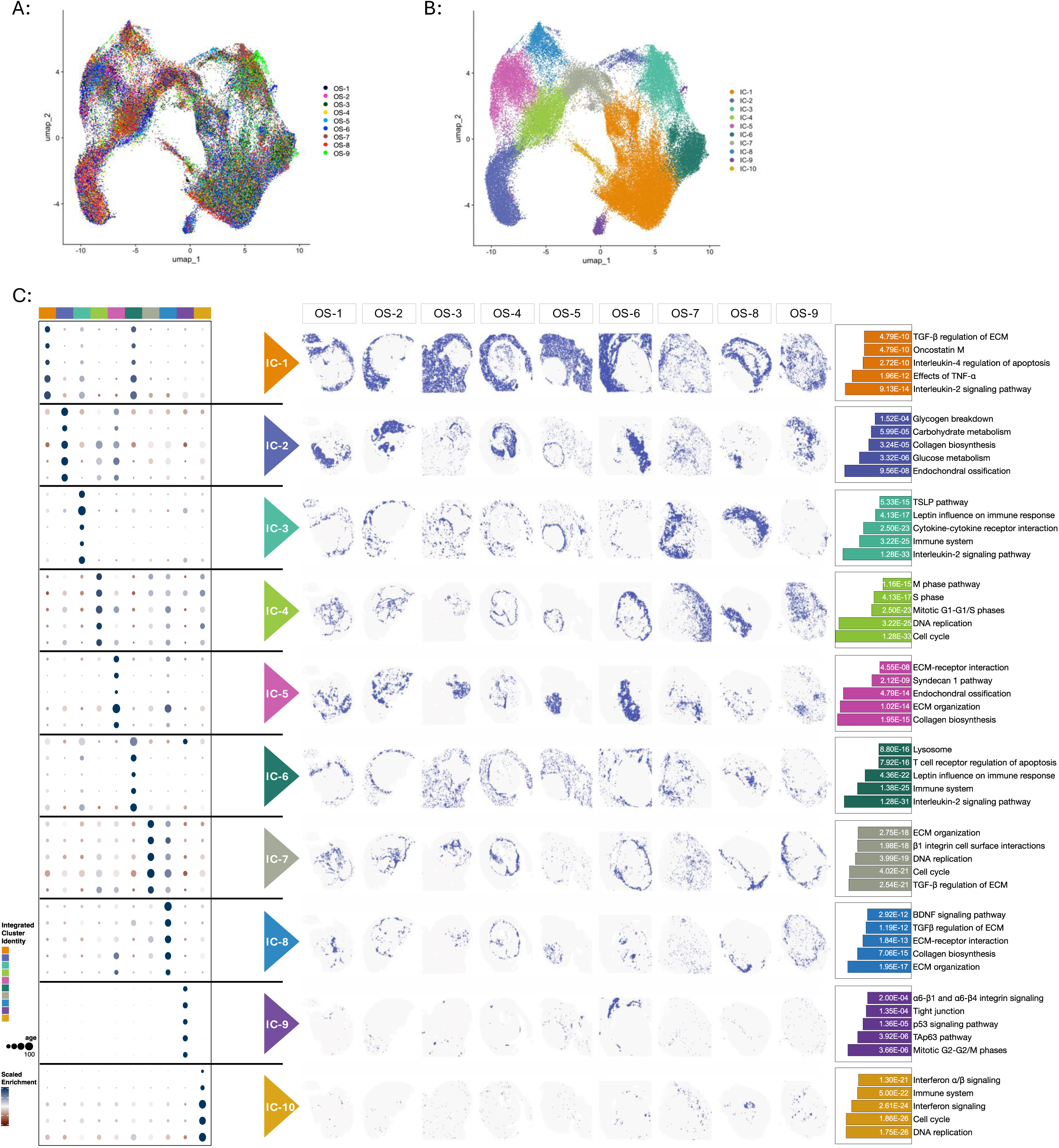
Metastatic osteosarcoma specimens exhibit a conserved transcriptional landscape. (A) UMAP of the integrated dataset after Harmony integration (theta=2, res=0.3, dims=15). Spots from individual specimens are represented in each cluster. (B) Cluster assignments of the integrated dataset. “IC” refers to Integrated Cluster. (C) Integrated clustering of metastatic osteosarcoma specimens. Left Panel: Dot-plot showing top 5 marker genes of each integrated cluster, designated as IC-1 through IC-10. Middle Panel: Spatial cluster maps localizing each integrated cluster within the tissues. A binary color palette was used to maximize contrast for purposes of display. Right Panel: Bar graphs of the most significantly enriched biological processes in each integrated cluster. P-values listed within each bar corresponds to biological process listed to the right.

### Drug-gene interaction analysis identifies CXCR4 as a clinically relevant therapeutic target

Next, we sought to leverage the conserved spatially resolved transcriptional architecture of metastatic osteosarcoma to identify region-specific druggable targets. Marker genes for each IC were used as input to identify drug-gene interactions in the Illuminating the Druggable Genome database. This query resulted in a list of Food and Drug Administration approved therapeutics targeting the druggable genes that define each IC. The drugs targeting each IC were largely unique with slight overlap between clusters (**Figure 7A**). Exceptions included IC-4 and IC-10, as well as IC-2 and IC-5, where considerable overlap in the drug lists were noted (**Figure 7A**). Of the drug-gene matches, the CXCR4 antagonist, plerixafor, had the highest score, preferentially targeting IC-3 (**Figure 7B**). Our analysis also showed that Plerixafor targets IC-5 and IC-6, albeit to a much lesser extent (data not shown). Importantly, of these IC’s targeted by plerixafor, IC-3 and IC-6 constitute the peritumoral microenvironment of each specimen which is hallmarked by a myeloid immune suppression transcriptional program (**Supplemental Figure 1**).^35,36^ To determine the clinical significance of these findings, we investigated the association between *CXCR4* expression and outcome of patients with osteosarcoma. This analysis demonstrated that *CXCR4* expression is moderately associated with inferior overall in patients with osteosarcoma **(Figure 7C)**. While the tumor tissue used to assess survival was obtained from sites of primary disease as opposed to to metastatic sites, these data indicate that CXCR4 is of potential clinical relevance and should be further investigated in the context of metastatic osteosarcoma.

**Figure 7:**
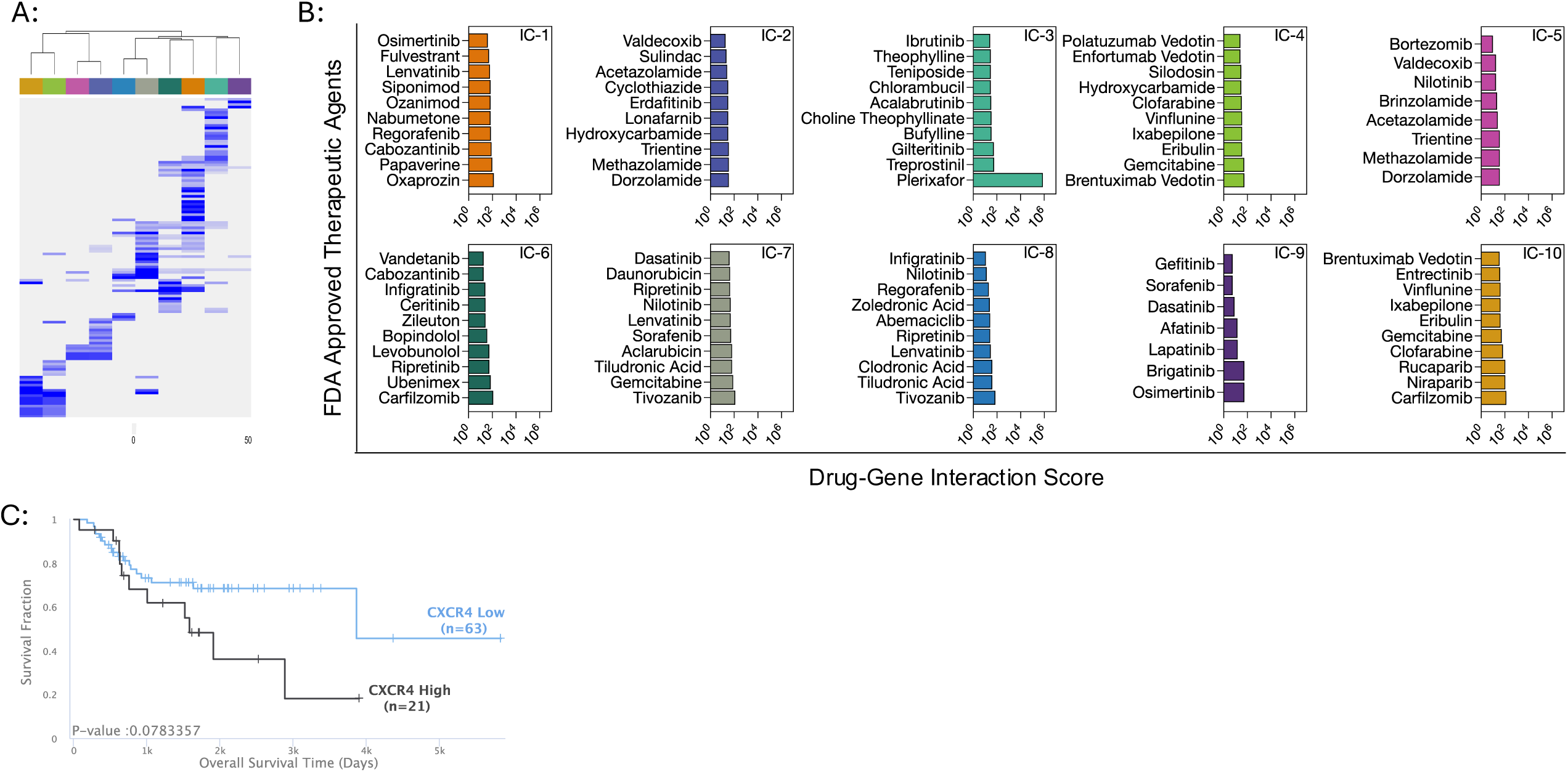
Identification and validation of the significance of the CXCR4 signaling axis as a potential therapeutic target. (A) Unsupervised hierarchical clustering of drug-gene interaction scores from the Illuminating the Druggable Genome database displayed as a heatmap. Rows indicate individual drugs. A higher score (blue) indicates a stronger interaction between a given drug and the query gene list. The color bar above the heatmap corresponds to the integrated spatial clusters. (B) Histograms of the top scoring drug-gene interactions for each integrated cluster. X-axis indicates drug-gene interaction score. (C) Kaplan-Meier plots showing the prognostic value of *CXCR4* expression in patients with osteosarcoma. P values < 0.05 are considered statistically significant.

### Regionally distinct expression patterns of CXCL12 and MIF suggests divergent CXCR4 signaling within the intratumoral and extratumoral microenvironments

CXCR4 is a G protein-coupled receptor whose ligands include CXCL12/SDF1 and MIF.^50^ To characterize the CXCR4 signaling axis in metastatic osteosarcoma, we examined the expression of *CXCR4, CXCL12,* and *MIF*. Spatial feature plots revealed widespread *CXCR4* expression with an enrichment localized to the peritumoral microenvironment (**Figure 8A**). Interestingly, we observed an inverse relationship between the expression patterns of *CXCL12* and *MIF* expression with *CXCL12* predominantly localized to the periphery and *MIF* expression concentrated within the intratumoral microenvironment (**Figure 8A**). These transcriptional data are consistent with our cytokine profiling results where MIF protein expression was readily detected in both tissue lysates obtained from specimens with high tumor cellularity as well as from conditioned media from patient-derived osteosarcoma cell lines cultured *in vitro* whereas CXCL12 protein expression was below the detection threshold (**Figure 3D-E**). Collectively, these data suggest that *MIF* is expressed by osteosarcoma cells and that *CXCL12* is expressed by peritumoral stromal and/or immune cells.

**Figure 8:**
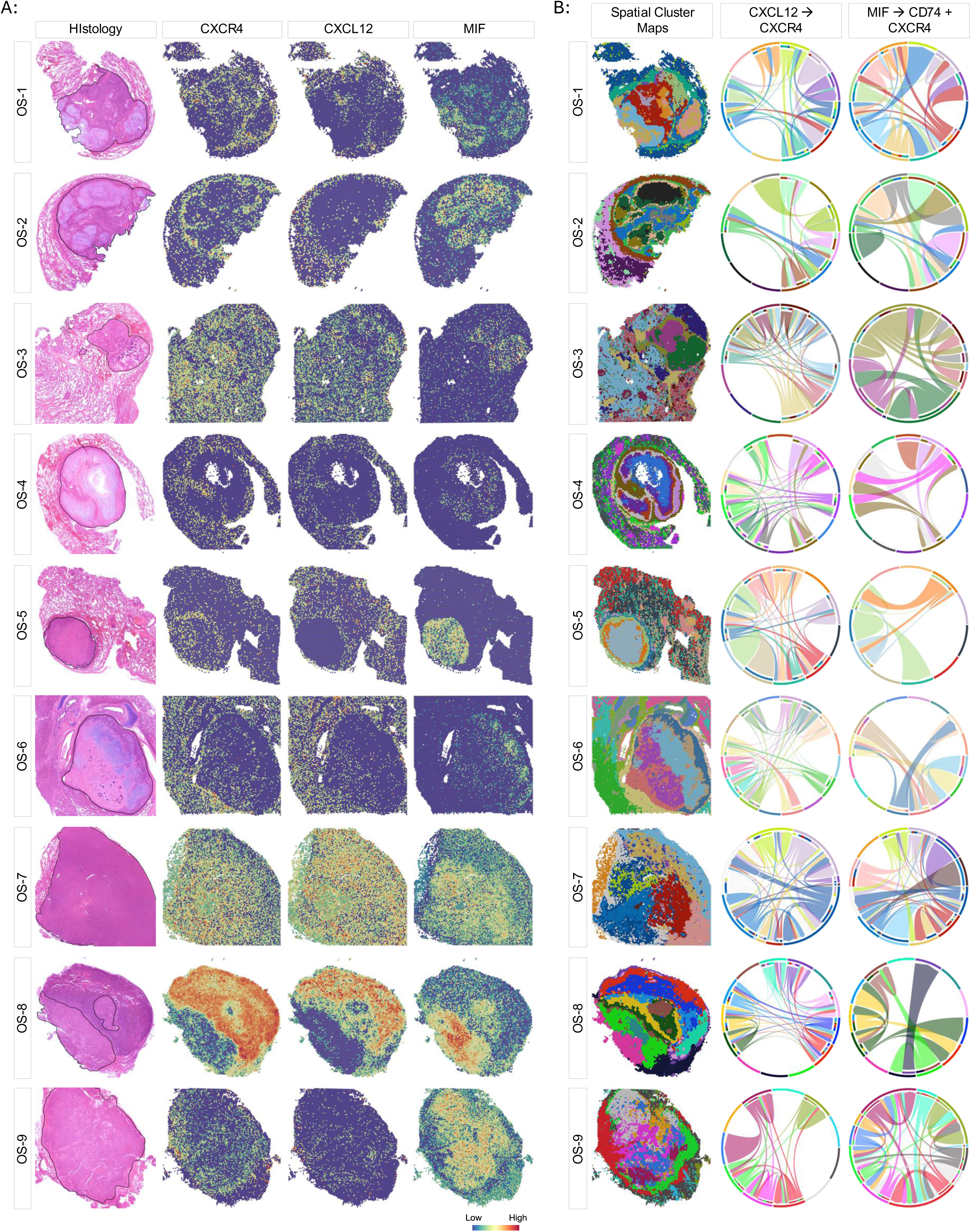
Regionally distinct CXCR4 ligand-receptor utilization. (A) Spatial gene expression plots showing the distribution of *CXCR4, CXCL12,* and *MIF* expression in each metastatic osteosarcoma specimen. Annotated hematoxylin and eosin-stained histological images are shown for reference. (B) Spatially resolved CXCR4 ligand-receptor interactome within and between spatially defined clusters. Spatial cluster maps are shown for reference. Interactome chord diagram is colored according to the spatial clusters displayed on the spatial cluster maps to the left.

Given the regional differences in ligand expression, we hypothesized that CXCR4 might engage different ligands in the intratumoral versus extratumoral microenvironment. To test this, we used the CellChat package to interrogate both canonical (CXCR4/CXCL12) and non-canonical (CXCR4/CD74/MIF) ligand-receptor interactions.^50^ This analysis uncovered spatial variability in ligand-receptor utilization, suggesting differential canonical and non-canonical CXCR4 signaling within and between the intratumoral and extratumoral microenvironments (**Figure 8B**). Although fibroblastic osteosarcoma specimen OS-7 did not exhibit the readily discernable spatial differences in *CXCR4, CXCL12,* and *MIF* expression observed in the other specimens (**Figure 8A**), ligand-receptor analysis did indeed provide evidence of regional differences in the CXCR4 signaling axis (**Figure 8B**). Taken together, these data highlight both the diversity of CXCR4 signaling and potential clinical utility in metastatic osteosarcoma. Detailed studies using genetically engineered and/or syngeneic murine models of metastatic osteosarcoma are necessary to experimentally elucidate the functional impact of CXCR4 signaling and its candidacy as a therapeutic target in this disease context.

## DISCUSSION

Exploring spatially-resolved cell-cell communication interactions within and between different microenvironments in metastatic osteosarcoma tissues can uncover therapeutic targets that are not evident from bulk genomic and/or transcriptomic approaches. Genomically, osteosarcomas are remarkably heterogenous tumors and due to this tremendous inter- and intra-tumoral genomic variability, osteosarcoma is often referred to as an “n=1” disease - a disease in which individual specimens are rarely alike.^51–53^ However, the extent to which this heterogeneity extends beyond the genome to the molecular composition and structure of these rare tumors has not been explored. Since osteosarcoma is a rare tumor obtaining fresh tissue from metastatic sites for detailed molecular profiling such as single cell and single nuclei sequencing, remains a challenge. Utilizing archived FFPE specimens to successfully perform spatial transcriptional profiling demonstrates the potential to provide unprecedented insight into the cellular and molecular composition of this aggressive disease. By using Visium slides with a 11mm x 11mm capture area, we were able to capture gene expression data from both the tumor and the surrounding non-malignant stromal tissue, allowing for a more comprehensive evaluation of how these diseased tissues are structured. Additionally, since this technique utilizes a single section as input, it is well suited for correlative biology studies to accompany investigational clinical trials, thereby providing robust datasets to enable more insight into the biologic impact of our therapies. The major limitations in our spatial molecular profiling study include the small sample size and that the 55mm resolution of the spots in the Visium assay used in this study which prohibits obtaining single cell level spatial gene expression data. This limitation hindered our ability to faithfully characterize individual subsets of intratumoral stromal cells and tumor cell populations as these gene signatures overlap. Importantly, by realizing this limitation we avoided overinterpretation of the data due to inaccurate assumptions.

Although a diagnosis of osteosarcoma necessitates the presence of malignant osteoid, these tumors are further classified as having osteoblastic, chondroblastic, or fibroblastic histology based on the preponderance of extracellular matrix observed.^26^ Including specimens of each histology in this profiling study enabled the evaluation of histological subtype as a variable. Of the specimens, fibroblastic osteosarcoma specimen OS-7 was the most divergent regarding tissue structure and transcriptional architecture, exhibiting a diffuse, rather than finely demarcated, tumor boundary. While entirety of the specimen, minus the lung parenchyma, was annotated as “tumor”, application of our Osteosarcoma Signature revealed a more nuanced tissue architecture with juxtaposing gradations of intratumoral and peritumoral microenvironments. This work demonstrates the utility of spatial transcriptional profiling to assist in refining the histopathological assessment of complex tumor specimens by identifying substructures not readily appreciated by histology alone.

We, and others, have previously shown that lymphocytes are excluded from infiltrating the intratumoral microenvironment of metastatic osteosarcoma.^25,27,54^ This, combined with the presence of peritumoral macrophages and multinucleated giant cells, circumscribing collagen deposition, and a peritumoral enrichment of the Defense Response signature, suggests that metastatic osteosarcoma invokes a foreign body-like response. As previously mentioned, osteosarcomas contain malignant osteoid, chondroid, and/or dense fibrous matrix with varying degrees of calcification.^26^ In the context of metastatic osteosarcoma (excluding bone metastases), these incredibly rigid tumors invade and grow in comparatively soft tissues. Based on the data presented here, we hypothesize that the rigidity and dense composition of the tumor and/or the development of ectopic malignant osteoid/chondroid damages the local tissues, thus initiating a foreign body-like/defense response, resulting in tumor encapsulation resemblant of a granuloma. As these metastatic lesions expand outward and induce additional local tissue damage, this response continues to propagate in a feed forward loop. We further hypothesize that the variability in the extent of this response reflects the different timing of metastatic seeding and outgrowth, resulting in differing amounts of matrix deposition in the fibrocollagenous capsule and severity/magnitude of the foreign body-like response. Our data suggests that this response is not limited to metastatic lesions within the lung parenchyma as we observe a similar response in the hilar lymph nodes as well. Although the lung is the most common site of osteosarcoma metastasis, spatial profiling of additional specimens from various anatomical disease sites will be necessary to determine the tissue type specificity of this response.

The purpose of a foreign body granuloma is to encapsulate the offending biologic or synthetic material to prevent further local tissue damage and systemic illness. Conceptually, the ability of metastatic lesions to provoke a foreign body-like response is presumably advantageous to the tumor by providing a physical barrier to protect it from destruction by the immune system. Mechanistically, the specific function of the metastatic encapsulation can be severalfold. Contracted collagen matrices have been shown to limit the diffusion of biomolecules.^55^ Our data demonstrates that there is minimal expression and activity of inflammatory chemokines and cytokines within the intratumoral microenvironment and that osteosarcoma cells themselves express few chemokines and cytokines. This is in stark contrast to the substantial inferred chemokine/cytokine activity in the extratumoral microenvironment that lies on the other side of the fibrocollagenous capsule. This coincides with the spatiotemporal dynamics of chemokine/cytokine signaling in chronic foreign body granulomas, whereby inflammatory signaling is absent within the core of the encapsulated lesions.^56,57^ It is possible that the encapsulation of metastatic lesions serves to restrain inflammatory signaling activity, providing a safe harbor for the tumor and tumor supporting cells to function and propagate.

Alternatively, extracellular matrix proteins have been shown to sequester chemokines and cytokines to direct signaling activities to a specific region.^58^ We have shown that the peritumoral microenvironment is consistently among the most actively communicating regions with regards to inflammatory chemokine/cytokine signaling. Moreover, macrophages are excellent cellular communicators that can orchestrate and transduce an array of multicellular processes.^30^ The distinct localization of immunosuppressive macrophages in the highly communicative peritumoral microenvironment, also supports a model whereby the dense circumscribing extracellular matrix functions to regionally concentrate immunological activity to modulate the tissue response to the presence of disease. Additional detailed *in vivo* experiments combined with high resolution single cell spatial profiling will be needed to truly determine the biological function of the peritumoral encapsulation of the metastatic lesions.

Patients with recurrent, refractory, and/or metastatic osteosarcoma have received limited clinical benefit from immune checkpoint blockade, yet the mechanisms underlying these clinical observations remain elusive.^59–62^ The observed regional imbalance of immunostimulatory versus immunosuppressive cytokines in the metastatic osteosarcoma tissues in this study hold important implications for checkpoint blockade and adoptive cell therapy. In both experimental murine models and patient specimens, TGFβ signaling mediates resistance to PD1/PD-L1 immune checkpoint blockade by promoting T-cell exclusion.^63^ By revealing widespread and heterogeneous TGFβ signaling activity, our data highlights the relevance of this pathway. However, the potential redundancy and/or compensatory signaling activity mediated by TGFβ, BMP, GDF, and Activin family members, illuminates the biological complexity of this pathway, warranting further mechanistic investigations to understand how to effectively target it.

We also observed high levels of SPP1, VEGFA, and CHI3L1 protein expression patient specimens. SPP1/osteopontin is a small integrin binding ligand N-glycosylated protein that is produced by various cell types within the osteosarcoma microenvironment including osteoblasts, osteocytes, fibroblasts, chondrocytes, myeloid-lineage cells, and endothelial cells.^64–67^ SPP1 promotes an immunosuppressive microenvironment via M2 macrophage polarization and suppression of CD8^+^ T-cells.^68,69^ Aside from its role in angiogenesis, VEGFA is a pleiotropic immunomodulatory cytokine that, when overexpressed in solid tumors, induces endothelial dysfunction, recruitment of M2 macrophages and myeloid derived suppressor cells, and promotes exhaustion of cytotoxic CD8^+^ T-cells.^70–77^ CHI3L1 is a glycoprotein that is upregulated in various tumor types and promotes immunosuppression by recruiting and activating tumor associated macrophages, myeloid derived suppressor cells, and regulatory T-cells.^78^ CHI3L1 also inhibits the activity of cytotoxic lymphocytes, induces fibrosis, and positively regulates TGFβ signaling.^78^

Our drug-gene interaction analysis identified plerixafor (Mozobil®) as a potential therapeutic molecule. Clinically, plerixafor is routinely used to mobilize hematopoietic stem cells in preparation for transplant by antagonizing CXCR4 on stem cells and thus allowing the cells to leave the bone marrow niche and enter the peripheral circulation. In addition to hematopoietic stem cells, CXCR4 is expressed on various cell types, including macrophages, myeloid derived suppressor cells, and osteosarcoma cells.^79–82^ Downstream CXCR4 signaling is incredibly complex with context dependent crosstalk with other receptors and ligand-specific downstream signal transducers and effectors.^80^ In its most simplistic form, CXCL12/SDF1 and MIF comprise the two major CXCR4 signaling axes. Activation of CXCR4 by CXCL12/SDF1 induces canonical downstream signaling whereas activation of CXCR4 by MIF and its coreceptor CD74, results in non-canonical signaling.^50,80^

In osteosarcoma cell lines, inhibition of the CXCL12/CXCR4 signaling axis reduced cell proliferation *in vitro* and attenuated metastatic progression *in vivo*.^79,81^ Similarly, pharmacologic inhibition of MIF reduced tumor cell growth and metastasis of osteosarcoma xenografts.^83^ Data exist showing that inhibition of CXCR4 signaling reduces intratumoral immunosuppression, resulting in increased efficacy of various immunotherapy regimens in murine models of breast cancer, pancreatic cancer, and melanoma.^84^ Although typically administered as a single subcutaneous injection, a phase I study examined the intratumoral effect of continuous infusion of plerixafor in patients with advanced pancreatic, ovarian, and colorectal cancers (NCT02179970). This study demonstrated that CXCR4 inhibition increased the abundance of tumor infiltrating lymphocytes into metastatic lesions from patients with microsatellite stable colorectal cancer and pancreatic ductal adenocarcinoma, sensitizing these tumors to checkpoint blockade.^85^ Currently, little is known about the functional impact of differential CXCR4 signaling in osteosarcoma and the subsequent effects on the structure and function of different spatial distinct microenvironments in metastatic disease. Nonetheless, the data from these studies provide intriguing rationale for CXCR4 signaling axis as a viable therapeutic target in metastatic osteosarcoma.

In conclusion, by shifting focus away from genomics and oncogenic mechanisms and more towards tissue structure and the spatial organization of cellular and molecular processes, our study provides new insight into the biology of pediatric metastatic osteosarcoma and the immunosuppressive mechanisms therein. Viewing metastatic disease through the perspective of a tumor associated foreign body-like response, our data shows that the transcriptional architecture and spatial landscape is conserved across metastatic osteosarcoma specimens, indicating a previously unappreciated commonality among disparate lesions, and challenging the prevailing “n=1” sentiment. Additional spatial profiling studies with a larger number of specimens will be needed to significantly increase our collective understanding of the regional specificity and interconnectedness of the various immunosuppressive mechanisms in osteosarcoma pulmonary metastases, thereby furthering our understanding of osteosarcoma disease biology. Elucidating how disseminated tumor cells alter the recipient tissue structure are paramount to revealing generalizable tissue-specific responses that can be further developed into innovative targeted therapies.

## ACKNOWLEDGEMENTS

This project has been funded by the Intramural Research Program of the National Cancer Institute of the National Institutes of Health (ZIABC012056 to TAM), the National Cancer Institute’s Childhood Cancer Data Initiative (3P30CA008748-54S3), and Federal funds under Contract No. 75N91019D00024. The Center for Pathology Research Services at Children’s Hospital Los Angeles and the Norris Comprehensive Cancer Center Translational Pathology Core are supported by grant 5P30CA014089-45. This work utilized the computational resources of the NIH High Performance Computing Biowulf cluster (http://hpc.nih.gov). The content of this publication does not necessarily reflect the views or policies of the Department of Health and Human Services, nor does mention of trade names, commercial products, or organizations imply endorsement by the United States Government.

## CONTRIBUTIONS

TAM conceptualized the study. MM, LK, JG, JS, and JC procured and evaluated patient specimens. JAB and JC provided histopathological expertise. PM, JP, DWM, CMS, JS, AC, MP, MK performed the experiments. JE, NW, PM, JAB, JCC, AKL, JC, RNK, and TAM analyzed and interpreted the data. TAM wrote the original manuscript. All authors reviewed and edited the manuscript.

---

**Supplemental Figure 1:**
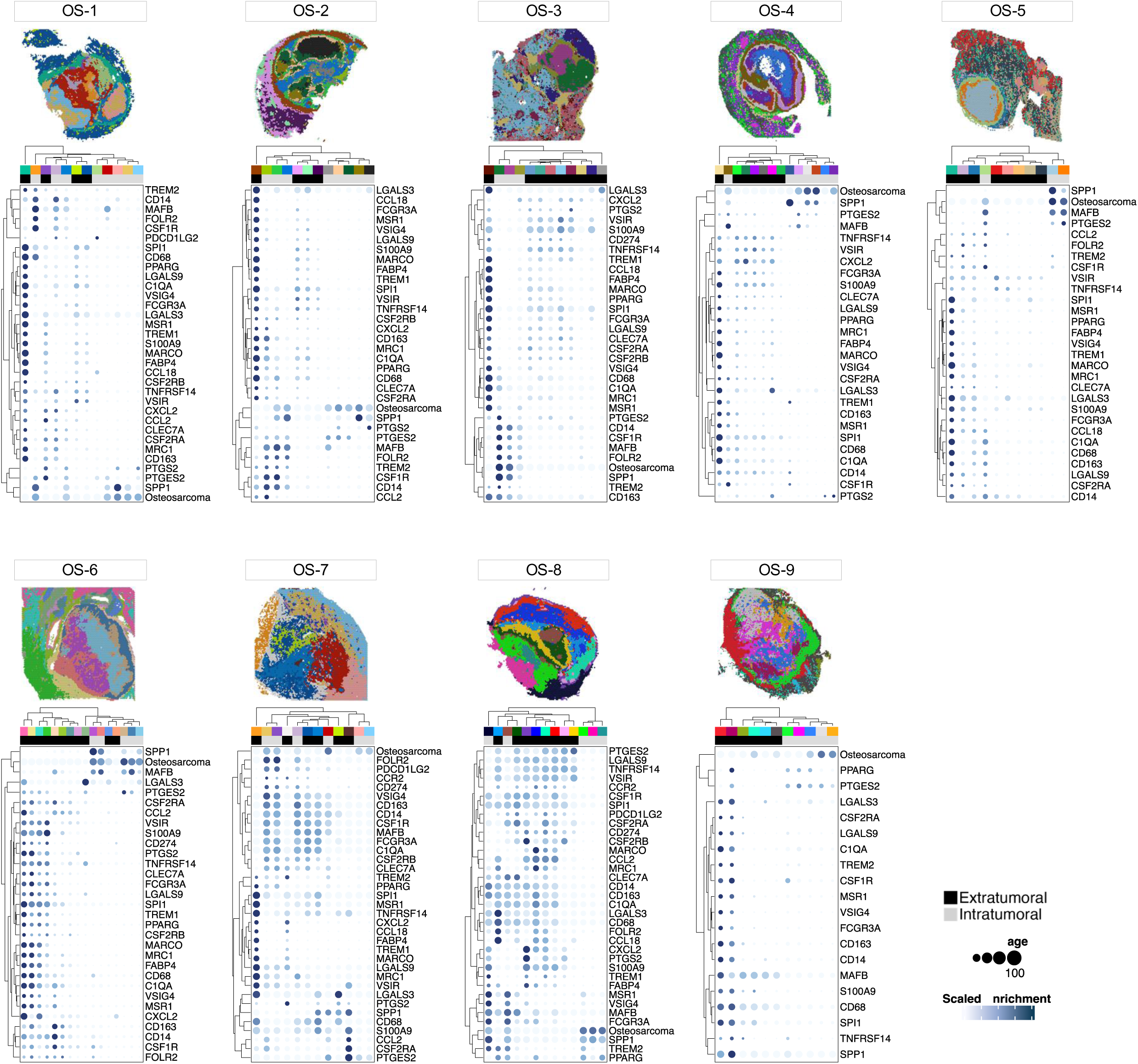
Distribution and abundance of genes encoding functional and phenotypic macrophage markers. Clustered dot plots illustrating the spatial distribution of gene encoding different functional and phenotypic macrophage markers within each individual metastatic osteosarcoma specimen. The color bar above each dot plot corresponds to the different spatial clusters displayed on the adjoining spatial cluster maps that are shown for reference (above). Separate color palettes denote the independent clustering analysis of each specimen. Spatial clusters were localized to the extratumoral (black) or intratumoral (gray) microenvironments as depicted in the second color bar.

**Supplemental Figure 2:**
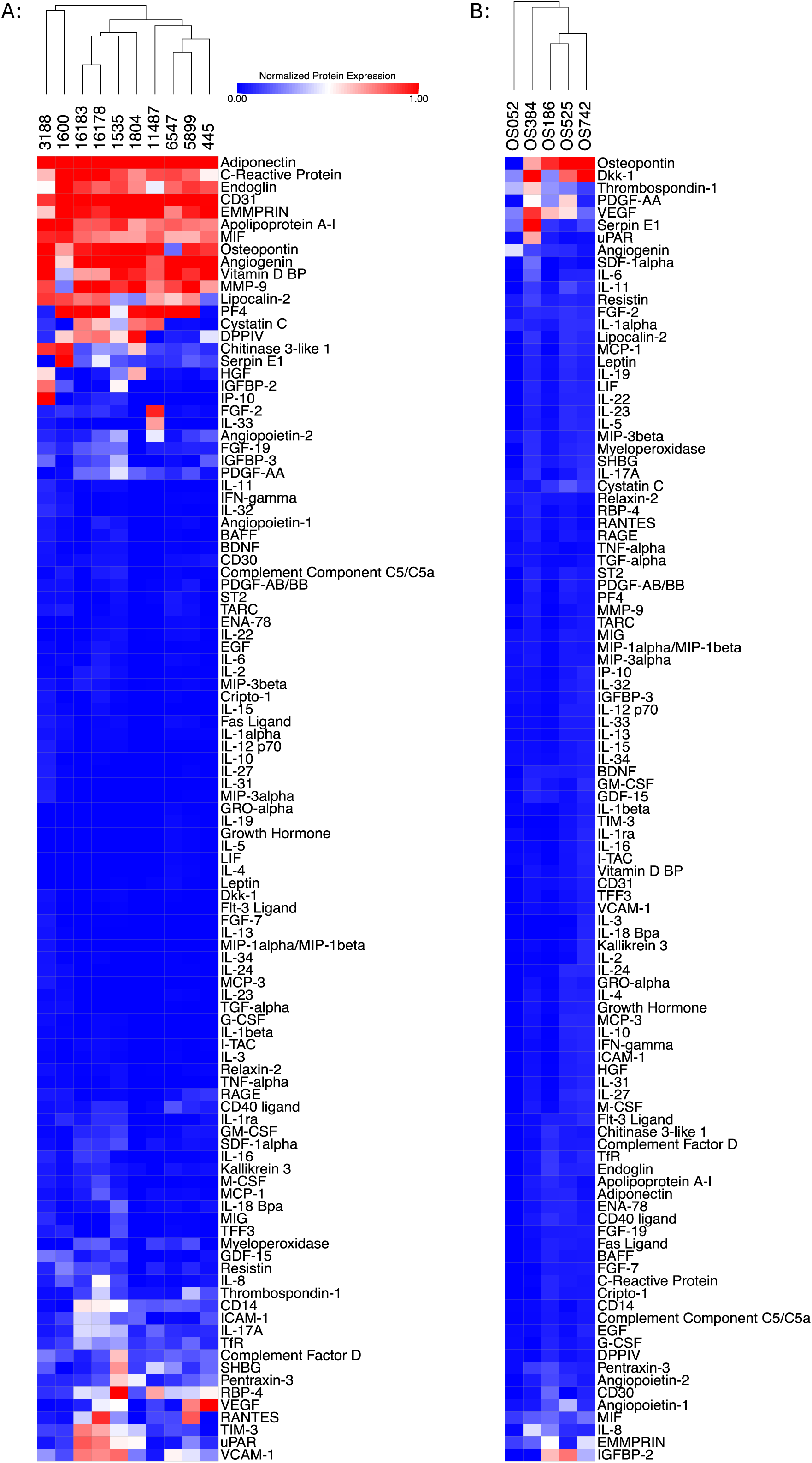
Chemokine/cytokine antibody array profiling in metastatic osteosarcoma patient specimens and patient derived cell lines. Heatmaps showing normalized protein expression of chemokines and cytokines from (A) metastatic osteosarcoma specimens and (B) patient derived cell line conditioned media.

**Supplemental Figure 3:**
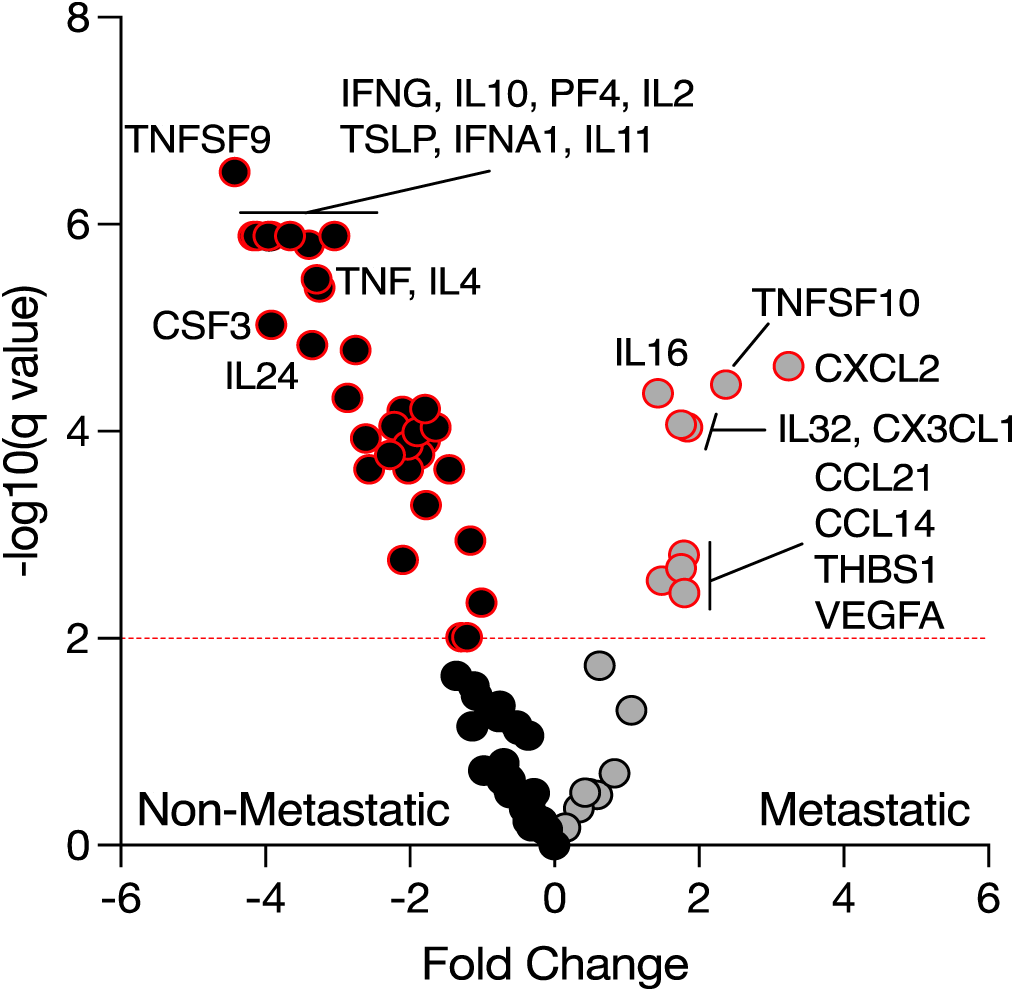
Differential gene expression of chemokine/cytokines in primary versus metastatic osteosarcoma specimens. Volcano plot of differentially expressed chemokine and cytokine genes in non-metastatic (black dots) and pulmonary metastases (gray dots) from the Sorenson, *et al.* NanoString IO360 gene expression dataset.^25^ Horizontal red line indicates significance threshold (q = 0.001). Dots with red outline indicates q < 0.001.

