## Supplemental Table 1 for "Spatial profiling identifies regionally distinct microenvironments and targetable immunosuppressive mechanisms in pediatric osteosarcoma pulmonary metastases"

| Long Name | Short Name | Genes |
|---|---|---|
| Adventitial Fibroblasts | Adventitial_Fibro | COL6A2, SFRP2, IGFBP6, MMP2 |
| Airway Smooth Muscle Cells | ASMC | DES, ACTA2, LGR6, DSTN, TPM2, TAGLN, MYH11, ACTG2, MYLK |
| Alveolar Epithelium Type 1 | AT1 | HOPX, AGER, RTKN2, EMP2, CLDN18, LMO7, CLIC3, KRT7 |
| Alveolar Epithelium Type 2 | AT2 | SFTPC, LAMP3, SLC34A2, SFTPB, SFTPA1, NPC2, NAPSA |
| Alveolar Fibroblasts type 1 | Alv_Fibro1 | TCF21, WNT2, SLC38A5, MEOX2, OLML3, VCAM1, COL13A1, ENPEP, ADH1B, G0S2, LBH, ITGA8, CDH11, PLXDC2, CDO1 |
| Alveolar Fibroblasts type 2 | Alv_Fibro2 | MFAP5, SCARA5, COL14A1, GPC3, CCDC80, RARRES2, LGALS1, PCOLCE, OGN, FSTL1 |
| Alveolar Macrophages | Alv_Mac | ALOX5AP, CD68, CTSD, FCER1G, MARCO, SPI1, SIGLEC1, ABCG1, FABP4 |
| Arterial Endothelial Cells | AEC | DKK2, GJA5, SERPINE2, HEY1, EFNB2, NOTCH1, BMX, CXCL12 |
| B Cells | B_cells | CD69, CORO1A, LIMD2, BANK1, LAPTM5, CXCR4, LTB, CD79A, CD37, MS4A1 |
| Basal Cells | Basal | KRT5, TP63, KRT14, NGFR, ITGA6, IGFBP4, ALCAM |
| Capillary Endothelial Cells Type 1 | CAP1 | APLNR, IL7R, GPIHBP1, FCN3, EDN1, SLC6A4, TEK |
| Capillary Endothelial Cells Type 2 | CAP2 | CA4, APLN, EDNRB, HPGD |
| CD4 T-cells | CD4_T | CORO1A, KLRB1, CD3E, LTB, CXCR4, IL7R, TRAC, IL32, CD2, CD3D |
| CD8 T-cells | CD8_T | CD8A, CD3E, CCL4, CD2, CXCR4, GZMA, NKG7, IL32, CD3D, CCL5 |
| Ciliated Airway Epithelium | Ciliated | FOXJ1, RSPH1, DYBLRB2, FAM183A, NME5, TPPP3, TUBA1A, TUBB4B, TMEM190 |
| Classical Monocytes | Class_Mono | LST1, IL1B, LYZ, COTL1, S100A9, VCAN, S100A8, S100A12, AIF1, FCN1 |
| Dendritic Cells | DC | CORO1A, MS4A6A, ITGB2, GPR183, HLA-DRB1, HLA-DPB1, HLA-DPA1, HLA-DQB1, HLA-DQA1, HLA-DMA |
| Fibromyocytes | Fibromyo | NEXN, ACTG2, LMOD1, PPP1R14A, DES, FLNA, TPM2, PLN, SELM |
| Inflammatory Monocytes | Infl_Mono | S100A8, S100A9, CD14, VCAN |
| Interstitial Macrophages | Int_Mac | C1QA, C1QB, C1QC, IL1B, MS4A4A, C3AR1, CD163, FCGR2A, NPL, SLCO2B1 |
| Ionocytes | Ionocyte | CFTR, FOXI1, ASCL3, ATP6V0B, AZGP1, HES6, TMEM61 |
| Lymphatic Endothelial Cells | LEC | PROX1, MMRN1, CCL21, PDPN, PTX3, NRP2, NRF2, FOXC2 |
| Mesothelial Cells | Meso | WT1, FREM2, UPK3B |
| Monocyte-derived Macrophages | Mono_Mac | LYZ, ACP5, TYROBP, LGALS1, CD68, AIF1, CTSL, EMP3, FCER1G, LAPTM5 |
| Myoepithelial Cells | MEC | KRT14, MYH11, TP63, KRT5, ACTA2, TAGLN |
| Myofibroblasts | Myofibro | CALD1, CYR61, TAGLN, MT1X, PRELP, TPM2, GPX3, CTGF, SPARCL1 |
| Neutrophils | Neut | S100A8, S100A9, IFITM2, FCGR3B, IL1B, CCR2, CSF1R |
| NK Cells | NK | GZMA, CD7, CCL4, CST7, NKG7, GNLY, CTSW, CCL5, GZMB, PRF1 |
| Non-Classical Monocytes | NonClass_Mono | PSAP, FCGR3A, FCN1, CORO1A, COTL1, FCER1G, LAPTM5, CTSS, AIF1, LST1 |
| Osteoclasts | Osteoclast | ACP5, CTSK, ITGB3, MMP13, MMP9 |
| Osteosarcoma Signature | Osteosarcoma | COL9A1, RUNX2, IBSP, MRC2, COL5A3, VIM, SPP1, MIK67, COL2A1, COL12A1, ALPL, LOX, TOP2A, COL11A1, COL11A2, PDGFD, CTHRC1, CLEC11A, TNC, COL27A1, SOX9, TGFBI, SATB2 |
| Patrolling Monocytes | Patrolling_Mon | CDKN1C, PTP4A3, HES4, TNFRSF8 |
| Pericytes | Pericyte | LAMC3, TRPC6, HIGD1B, PDGFRB, COX4I2, KCNK3, NOTCH3, CSPG4, |
| Plasma Cells | Plasma | ITM2C, TNFRSF17, FKBP11, IGKC, IGHA1, IGHG1, CD79A, JCHAIN, MZB1, ISG20 |
| Plasmacytoid Dendritic Cells | pDC | CLEC4C, LILRA4, IRF7, PLD4 |
| Regulatory T-cells | Treg | FOXP3, CTLA4, IL2RA |
| Serous Cells | Serous | LTF, LYZ, CCL28, SAA2, PIGR, SLPI |
| Smooth Muscle Cells | SMC | PRKCDBP, NDUFA4L2, MYL9, ACTA2, MGP, CALD1, TPM1, TAGLN, IGFBP7, TPM2 |
| Subpleural Fibroblasts | Subpleural_Fibro | SERPING1, C1R, NNMT, MT1E, MT1X, PLA2G2A, SELM, MT1M |
| Systemic Venous Endothelial Cells | SVEC | ACKR1, COL15A1, ABCB1, VWA1 |
| Vascular Endothelial Cells | VEC | ACKR1, PRDD23, VWF, CLU, EPHB4, IGFBP7, NR2F2 |

Supplemental Table 1
