## Supplemental Table 2 for "Spatial profiling identifies regionally distinct microenvironments and targetable immunosuppressive mechanisms in pediatric osteosarcoma pulmonary metastases"

| Target | Tag | Clone | Dilution | Catalog ID | Vendor |
|---|---|---|---|---|---|
| A-SMA | 141Pr | 1A4 | 600 | 3141017D | SBT |
| Vimentin | 143Nd | D21H3 | 300 | 3143027D | SBT |
| CD68 | 159Tb | KP1 | 300 | 3159035D | SBT |
| CD8a | 162Dy | D8A8Y | 200 | 3162035D | SBT |

Supplemental Table 2
